# *Salmonella enterica* serovar Typhimurium from Wild Birds in the United States Represent Distinct Lineages Defined by Bird Type

**DOI:** 10.1101/2021.11.26.470158

**Authors:** Yezhi Fu, Nkuchia M. M’ikanatha, Chris A. Whitehouse, Shaoting Li, Xiangyu Deng, Jared C. Smith, Nikki W. Shariat, Erin M. Nawrocki, Edward G. Dudley

## Abstract

*Salmonella enterica* serovar Typhimurium is typically considered a host generalist, however certain strains are associated with specific hosts and show genetic features of host adaptation. Here, we sequenced 131 *S.* Typhimurium strains from wild birds collected in 30 U.S. states during 1978-2019. We found that isolates from broad taxonomic host groups including passerine birds, water birds (Aequornithes), and larids (gulls and terns) represented three distinct lineages and certain *S.* Typhimurium CRISPR types presented in individual lineages. We also showed that lineages formed by wild bird isolates differed from most strains originating from domestic animal sources, and genomes from these lineages substantially improved source attribution of Typhimurium genomes to wild birds by a machine learning classifier. Furthermore, virulence gene signatures that differentiated *S.* Typhimurium from passerines, water birds, and larids were detected. Passerine isolates tended to lack *S.* Typhimurium-specific virulence plasmids. Isolates from the passerine, water bird, and larid lineages had close genetic relatedness with human clinical isolates, including those from a 2021 U.S. outbreak linked to passerine birds. These observations indicate that *S.* Typhimurium from wild birds in the U.S. are likely host-adapted, and the representative genomic dataset examined in this study can improve source prediction and facilitate outbreak investigation.

**IMPORTANCE:** Within-host evolution of *S.* Typhimurium may lead to pathovars adapted to specific hosts. Here, we report the emergence of disparate avian *S.* Typhimurium lineages with distinct virulence gene signatures. The findings highlight the importance of wild birds as a reservoir for *S.* Typhimurium and contribute to our understanding of the genetic diversity of *S.* Typhimurium from wild birds. Our study indicates that *S.* Typhimurium may have undergone adaptive evolution within wild birds in the U.S. The representative *S.* Typhimurium genomes from wild birds, together with the virulence gene signatures identified in these bird isolates, are valuable for *S.* Typhimurium source attribution and epidemiological surveillance.

## INTRODUCTION

*Salmonella enterica* is a major zoonotic and foodborne pathogen, and one of the leading bacterial causes of foodborne illnesses in the U.S. The Centers for Disease Control and Prevention (CDC) estimates that *Salmonella enterica* is responsible for approximately 1.35 million illnesses, 26,500 hospitalizations, and 420 deaths each year in the U.S.(1). Food products such as eggs, livestock, and poultry meat are the main sources for most of these illnesses (1). Source attribution studies have documented that food-producing animals such as pigs, cattle, and chicken are the main reservoirs for *Salmonella* bacteria that infect humans (2). Although transmission of non-typhoidal *Salmonella* to humans through food and food-producing animals is well described, the contribution of nonfood sources such as wild birds is less defined. Nonfood-producing animals are less explored in most source-attribution studies mainly due to a lack of representative and routinely collected data on these animals (3).

There have been several reports suggesting transmission of *S. enterica* between wild birds and humans in recent years. For instance, possible interspecies transmission of *S.* Enteritidis among gulls, poultry, and humans has been reported in Chile (4). Isolates of *S.* Hvittingfoss from a 2016 multi-state outbreak originating from tainted cantaloupes in Australia closely matched *S.* Hvittingfoss strains isolated from bar-tailed godwits (5). Although there is no direct evidence suggesting that the above-mentioned outbreaks originated from charadriiform birds (shorebirds and larids), it highlights the potential role of charadriiform birds in dissemination of *S. enterica* involved in human infections. Investigations focused on passerine birds (songbirds) identified a total of 337 human infections with passerine-associated *S.* Typhimurium DT40, DT56(v) and DT160 isolates during 2000-2010 (6). In 2021, an *S.* Typhimurium outbreak linked to passerines has led to 29 illnesses and 14 hospitalizations in over 12 U.S. states (7).

As wild birds may serve as a reservoir of *S. enterica* that cause human salmonellosis, it is important to characterize isolates from wild birds so that strains from human cases can be properly traced to the appropriate source and potential transmission routes can be identified. A public portal integrating pathogen genomic sequences from various sources and geographic regions can facilitate source attribution studies with large datasets in real-time. Although the NCBI Pathogen Detection (PD) provides such a data portal for traceback investigation of an outbreak (https://www.ncbi.nlm.nih.gov/pathogens/), the sequencing data are skewed towards isolates from clinical samples, foods, or food-producing animals (i.e., livestock and poultry). The limited data from nonfood sources such as wild birds may hinder source attribution during an outbreak investigation, thus affecting subsequent prevention and mitigation measures. Besides direct sequence comparisons, machine learning algorithms such as Random Forest (RF) classifier have shown promise for predicting *Salmonella* host specificity (8) or zoonotic sources (9). Zhang et al. reported a Random Forest classifier for genomic source prediction of *S.* Typhimurium in the U.S. that showed a marked difference in prediction accuracies between food-producing animals (prediction accuracy ~90%) and wild birds (prediction accuracy ~50%). The difference was presumably due to a lack of representative and available wild bird isolates (9). Therefore, it is necessary to include *S. enterica* isolates beyond food or food-producing animals in a public data portal such as PD to help better resolve complex *Salmonella* epidemiology.

In this study, we sequenced 131 *S.* Typhimurium strains from wild birds isolated by the U.S. Geological Survey - National Wildlife Health Center. Strains were isolated between 1978-2019 and originated from birds collected in 30 U.S. states. We used these data to test the following hypotheses: 1) Strains would be phylogenetically segregated by bird host; 2) Additional *S*. Typhimurium genomes from wild bird isolates would improve the performance of machine learning-based source attribution; 3) Allelic variation in virulence genes would contribute to *S.* Typhimurium host specificity; and 4) Inclusion of these data in the PD database would better define cross-species transmission of *S.* Typhimurium between wild birds, humans, and other hosts.

## RESULTS

### Distinct *S.* Typhimurium lineages are carried by passerine birds, water birds, and larids

Metadata (isolate name, SRA accession number, collection year, state, source/host species, CRISPR type, and classic 7-gene MLST sequence type) for the 131 isolates are listed in Table S1. Based on single nucleotide polymorphism (SNP) phylogenetic analyses, isolates from wild birds formed three major lineages (**Fig. 1A**), each supported by robust bootstrap values of 100%. The largest branch included 42 isolates, all of which originated from passerine birds (order Passeriformes) (e.g., cardinal, finches, and sparrows). The second major lineage included 40 isolates predominantly originating from water birds of the clade Aequornithes (e.g., cormorants, pelicans, and herons). This lineage also included occasional strains isolated from other birds that commonly share the same aquatic habitats or prey/scavenge on the carcasses of the above-mentioned species: terns (*n* = 2), gull (*n* = 1), grebe (*n* = 1), goose (*n* = 1), and bald eagle (*n* = 1). The third major *S.* Typhimurium lineage consisted of 32 strains primarily isolated from larids (terns and gulls; order Charadriiformes). This lineage also included isolates originating from a plover (also a member of Charadriiformes) and an owl. Subclades formed by *S.* Typhimurium strains within each of the three major lineages did not show clear patterns of host specificity (e.g., strains isolated from terns and gulls did not form unique subclades but rather were interspersed within the core larid lineage). Isolates from raptors (e.g., owls, bald eagles, and hawks) did not cluster together, and are highlighted in red in **Fig. 1A**. Based on Bayesian phylogenetic inference (**Fig. 1B**), we were able to estimate that the most recent common ancestor (MRCA) of the water bird lineage was from around 1915 [95% highest posterior distribution (HPD): 1902-1926]. The MRCAs of the passerine lineage and larid lineage were from approximately 1941 (95% HPD: 1934-1947) and 1938 (95% HPD: 1929-1947), respectively. The passerine lineage and the larid lineage were estimated to have split from each other around 162 years ago (median: 1857; 95% HPD: 1841-1873).

**FIG 1.**
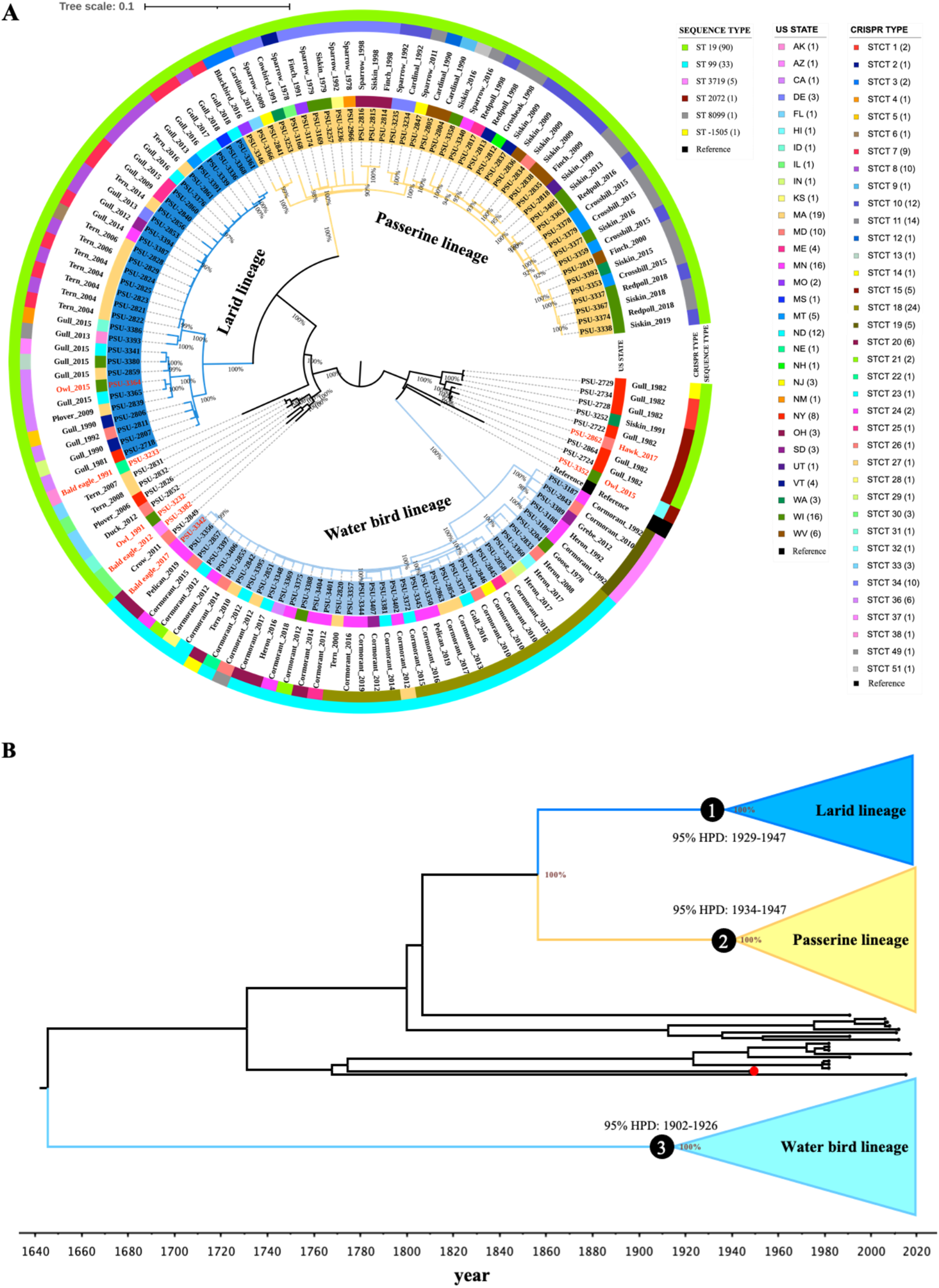
Phylogenetic analyses reveal distinct *S.* Typhimurium lineages in passerine birds, water birds, and larids. (A) Maximum-likelihood phylogenetic tree of 131 *S.* Typhimurium isolates from wild birds collected between 1978-2019 in 30 U.S. states. The tree is visualized using iTOL. Three large lineages are defined by bird types, i.e., passerine lineage (yellow), larid lineage (blue), and water bird lineage (light blue). Bootstrap values are displayed as percentage on tree branch. The labels at tips represent the isolate name, the inner color strip represents the U.S. state from which the sample originated, the outmost color strips represent the *S.* Typhimurium CRISPR type and MLST sequence type, and the text labels between the color strips represent the bird host and isolation year. The legend fields at the right of the tree represent the *S.* Typhimurium MLST sequence type (number of isolates), U.S. state name (number of isolates), and the *S.* Typhimurium CRISPR type (number of isolates). The labels highlighted in red represent isolates from raptors. (B) Time-scaled Bayesian phylogenetic tree of 131 *S.* Typhimurium isolates from wild birds collected between 1978-2019 in 30 U.S. states. The tree is visualized using FigTree v1.4.4. Numbers in black circles at the nodes represent the 95% highest posterior probability density (HPD) for the times of most recent common ancestor for the water bird lineage (light blue), larid lineage (blue), and passerine lineage (yellow). The red circle at the tree tip represents the reference strain LT2 (isolation year: 1948). Posterior probability values are displayed as percentage on tree nodes of the three lineages.

### Defined *S.* Typhimurium lineages contain disparate CRISPR types

We further classified isolates by two established *Salmonella* typing methods, i.e., classic 7-gene multilocus sequence typing (MLST) and CRISPR (Clustered Regularly Interspaced Short Palindromic Repeats) typing, to determine the sequence type (ST) and *S.* Typhimurium CRISPR type (STCT) of our isolates. Six STs were detected and 93.9% of the isolates (123/131) belonged to ST19 (*n* = 90) and ST99 (*n* = 33). All the ST99 isolates belonged to the water bird lineage. However, ST did not always distinguish between the lineages. Specifically, ST19 isolates were dispersed among the larid and passerine bird lineages (**Fig. 1A** and **Table S1**). On the other hand, CRISPR analysis identified 37 STCTs within the 131 isolates (**Fig. 1A** and **Table S1**), several of which only appeared in certain lineages. For examples, STCT10 (*n* = 12), STCT11 (*n* = 14), and STCT34 (*n* = 10) were unique to the passerine bird lineage; STCT18 (*n* = 24), STCT19 (*n* = 5), and STCT20 (*n* = 6) only occurred in the water bird lineage; and STCT7 (*n* = 9), STCT8 (*n* = 10), and STCT36 (*n* = 6) were exclusive to the larid lineage.

### Genetic relatedness of U.S. wild bird isolates with isolates from other reservoirs and geographic regions

We examined whether wild bird isolates of *S*. Typhimurium were related to those from other host sources and geographic regions. **Fig. 2** and **Table S2** show that wild bird isolates were related to other isolates in the PD database from diverse sources (e.g., humans, water, fish/shellfish, horses, cats, and food) and different geographic regions. The PD SNP Tree Viewer showed that our isolates from the passerine lineage, water bird lineage, and larid lineage formed six, four, and four SNP clusters with other isolates in the large database, respectively (**Table S2**). Each SNP cluster contains isolates whose genomes are within 50-SNP distance of each other. We summarized the source niches and geographic regions of all the isolates in the SNP clusters formed by different *S.* Typhimurium lineages, and the results are represented in **Fig. 2**. As shown in **Fig. 2, A** and **B**, isolates from both the water bird and larid lineages had close genetic relatedness to isolates from water, fish/shellfish, etc. In contrast, isolates of the passerine lineage were not related to fish/shellfish isolates in the PD database; rather these isolates were more closely related with strains originating from horses and cats (**Fig. 2C**). Notably, isolates from the three major wild bird lineages were genetically similar to a number of clinical isolates from humans. The numbers of clinical isolates clustered with isolates from water bird, larid, and passerine lineages in the PD database were 37, 84, and 495, respectively. Interestingly, isolates from food-producing animals such as livestock and poultry rarely clustered with wild bird isolates. Only two isolates from livestock were related to passerine-derived isolates, while none was closely associated with water bird or larid isolates (**Fig. 2, A** to **C**). Geographically, isolates in both the water bird and larid lineages were closely related to strains originating from outside of the U.S. Isolates from Canada, Chile, Mexico, UK, Japan, and Australia clustered with isolates from the water bird and larid lineages from the U.S. (**Fig. 2, D** and **E**). Isolates from the passerine lineage only demonstrated close genetic relatedness with strains isolated from sources within North America (**Fig. 2F**).

**FIG 2.**
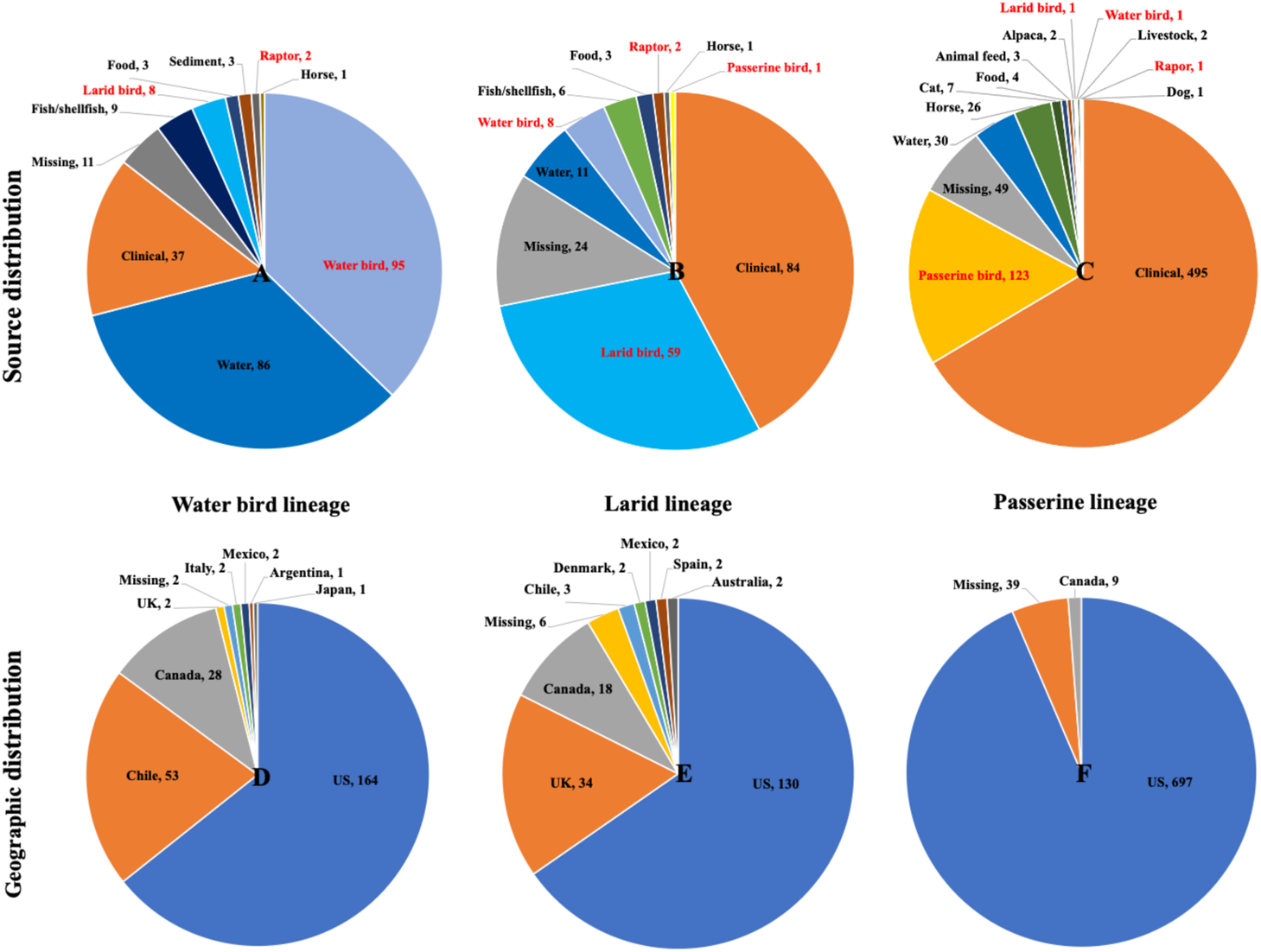
Distribution of source niches and geographic regions for the isolates in the Pathogen Detection (PD) database that cluster with *S*. Typhimurium isolates from different wild bird lineages. (A-C) Source distribution of isolates in the PD database clustering with isolates from water bird lineage, larid lineage, and passerine lineage. (D-F) Geographic distribution of isolates in the PD database clustering with isolates from water bird lineage, larid lineage, and passerine lineage. The number following the source niche or geographic region represents the number of isolates from that specific source or geographic region in the PD database. Complete data used to generate the pie charts are given in Table S2.

### Inclusion of the 131 *S.* Typhimurium isolates from wild birds improves source prediction accuracy of a Random Forest classifier

Given the distinct clustering of wild bird isolates, we next explored whether our dataset would improve a previously published Random Forest (RF) classifier for source prediction of *S*. Typhimurium strains. With the original training dataset (195 bovine isolates, 338 porcine isolates, 440 poultry isolates, and 68 wild bird isolates), the RF classifier had a prediction accuracy of 64.62%, 88.76%, 91.14%, and 52.94% for bovine, porcine, poultry, and wild birds, respectively, and an out-of-bag prediction accuracy of 82.9%. After adding our 131 wild bird isolates into the training dataset, the out-of-bag prediction accuracy was increased to 84.3% (**Table 1**). Notably, there was a substantial increase in prediction accuracy for wild birds from 52.94% to 83.42%. In contrast, the prediction accuracies for bovine and poultry remained largely unaffected. To explain the prediction results using phylogeny, a SNP tree was built that included 1,604 isolates from different sources [171 human clinical isolates, 261 isolates from food and other sources, and 1,172 isolates used for the RF classifier training (i.e., 195 bovine isolates, 338 porcine isolates, 440 poultry isolates, and 199 wild bird isolates in which 131 were from this study)]. These different sourced isolates excluding the 131 wild bird isolates were characterized in a previous *S.* Typhimurium source attribution study (9) The wild bird isolates formed three lineages that did not overlap with those formed by other animal-sourced isolates (**Fig. 3**), which corresponded to the source prediction performance of the RF classifier that the number of wild bird isolates falsely attributed to porcine, bovine, or poultry sources only increased from 32 to 33 after adding our bird isolates into the training dataset (**Table 1**). It should be noted that isolates from bovines were more dispersed and overlapped with other sourced isolates in the phylogenetic tree (**Fig. 3**). This was consistent with the source prediction performance of the RF classifier that 37.44% (73/195) of the bovine isolates were assigned to porcine, poultry, or wild bird sources (**Table 1**).

**FIG 3.**
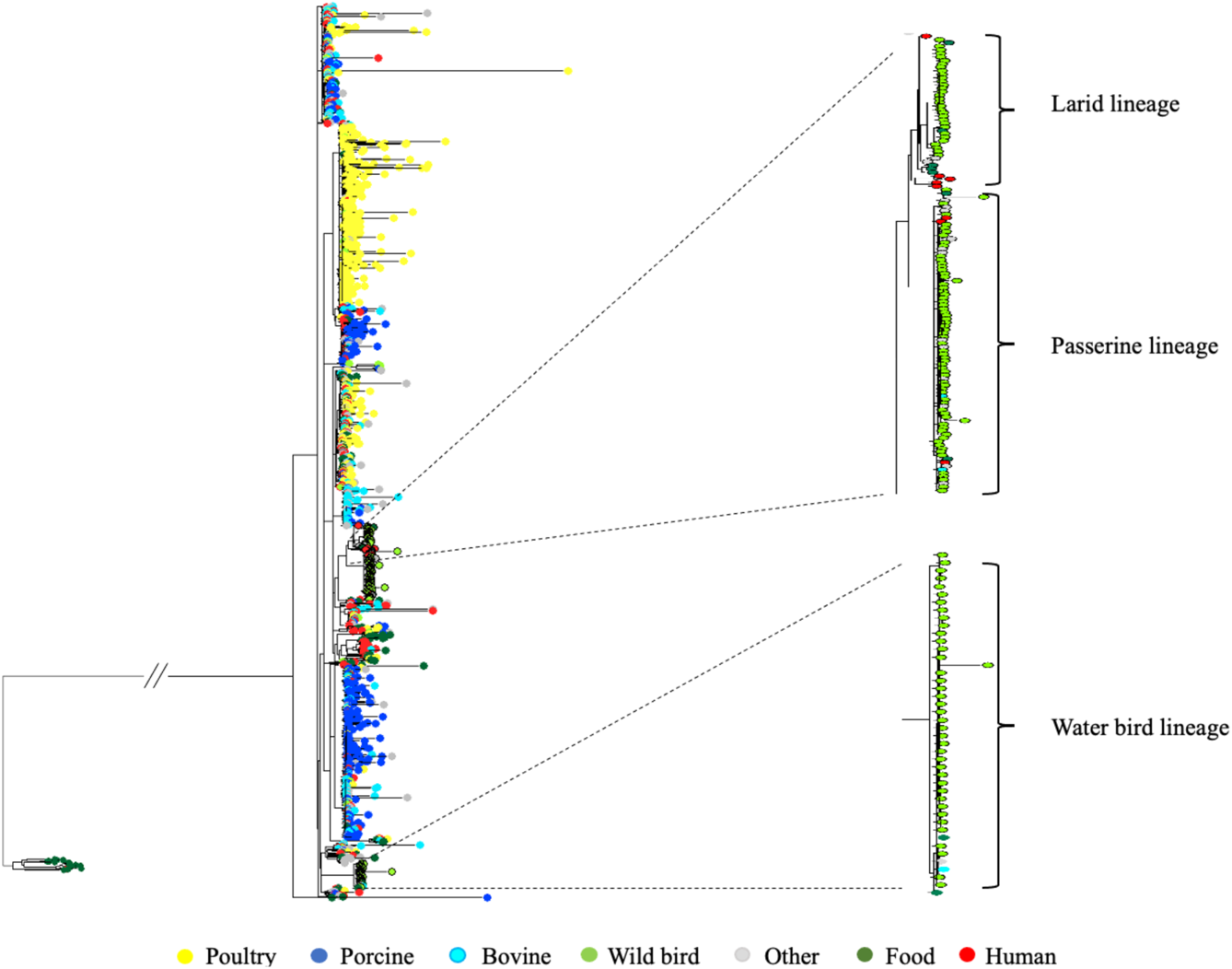
Maximum-likelihood phylogenetic tree of 1,604 S. Typhimurium isolates from different sources for zoonotic source prediction. The maximum-likelihood phylogenetic tree for zoonotic source prediction includes S. Typhimurium isolates from seven source classes: (1) bovine (n = 195), including isolates from cattle, beef, and raw milk; (2) porcine (n = 338), including isolates from pigs and pork; (3) poultry (n = 440), including isolates from chickens, turkeys, ducks, and their eggs; (4) wild bird (n = 199), including 131 isolates from this study; (5) human (n = 171), including human clinical isolates; (6) food (n = 83), including seafood, plant-based food, and other ready-to-eat and/or processed food; (7) other (n = 178), including any isolates not belonging to the aforementioned classes. The larid, passerine, and water bird lineages formed by the S. Typhimurium isolates from wild birds are enlarged and highlighted in the phylogenetic tree.

**Table 1.** Confusion matrix showing the prediction accuracy of a Random Forest classifier for source attribution of *S*. Typhimurium to bovine, porcine, poultry, and wild bird sources

| **Original training dataset** |  |  |  |  |  |
| --- | --- | --- | --- | --- | --- |
| Out-of-bag prediction accuracy: 82.9% |  |  |  |  |  |
| Predicted class<br>Actual class | Bovine | Porcine | Poultry | Wild bird | Class accuracy |
| Bovine | 126 | 38 | 29 | 2 | 64.62% |
| Porcine | 24 | 300 | 10 | 4 | 88.76% |
| Poultry | 24 | 11 | 401 | 4 | 91.14% |
| Wild bird | 10 | 8 | 14 | 36 | 52.94% |
| **Original training dataset with addition of the 131 wild bird isolates** |  |  |  |  |  |
| Out-of-bag prediction accuracy: 84.3% |  |  |  |  |  |
| Predicted class<br>Actual class | Bovine | Porcine | Poultry | Wild bird | Class accuracy |
| Bovine | 122 | 41 | 28 | 4 | 62.56% |
| Porcine | 26 | 300 | 8 | 4 | 88.76% |
| Poultry | 24 | 12 | 400 | 4 | 90.91% |
| Wild bird | 12 | 7 | 14 | 166 | 83.42% |

### Genetic signatures related to virulence differ between strains originating from different types of wild birds

We next took a more focused approach to examine whether presence of antimicrobial resistance genes, virulence genes and/or plasmids were associated with the major wild bird lineages. All 131 wild bird *S*. Typhimurium strains were predicted to be sensitive to major types of antimicrobials (beta-lactam, tetracycline, phenicol, fluoroquinolone, etc.) (**Table S3**). These isolates carried common virulence determinants found in *S.* Typhimurium. These included fimbrial/nonfimbrial adherence determinants, genes promoting bacterial survival in activated macrophage, type three secretion system (TTSS) encoded by *Salmonella* pathogenicity island I and II (SPI-1 and SPI-2), and genes associated with magnesium uptake, serum resistance, stress adaptation, and toxin production (**Table S3**).

Isolates from the water bird, larid, and passerine lineages each had defining mutations that may abolish the function of specific virulence genes. The *fimC* gene (full length: 708 bp) encoding the type 1 pilus chaperone FimC had a single base-pair deletion at position 87 in all isolates (100%; 42/42) belonging to the passerine lineage. Only two isolates (i.e., PSU-2849 and PSU-3252) from passerine birds were predicted to encode the full length *fimC*, however these isolates were divergent from the core passerine lineage. Conversely, all the isolates (100%; 72/72) from the water bird and larid lineages possessed an intact *fimC* (**Fig. 4A**). On the other hand, all the isolates (100%; 40/40) in the water bird lineage had a single base-pair insertion in the *safB* gene (full length: 714 bp; insertion at position 601) encoding the Saf pilus chaperone SafB, a 60 base-pair deletion in the *sthC* gene (full length: 2538 bp; deletion at 481-540) encoding the Sth pilus usher protein SthC, and a single base-pair insertion and substitution in the *sseL* gene (*<u>S</u>almonella* <u>s</u>ecreted <u>e</u>ffector <u>L</u>; full length: 962 bp; insertion at position 474, substitution at position 823). These three genes occurred in all the isolates from passerine and larid lineages (**Fig. 4A**). However, there was a substitution at position 867 of the *sseL* gene in isolates from the larid lineage that led to a premature stop codon (**Fig. S1**). The organization of the fimbrial operons containing *fimC*, *safB*, and *sthC* in isolates from the three major wild bird lineages, and reference isolate *S.* Typhimurium LT2 are shown in **Fig. 4, B** to **D**. Gene *sseL* was monocistronic and its alignment within different bird isolates is shown in Fig. S1. Overall, the mutations (frameshift and premature stop codon) of these virulence genes resulted in pseudogene accumulation that may inactivate their functions (**Fig. 4, B** to **D**).

**FIG 4.**
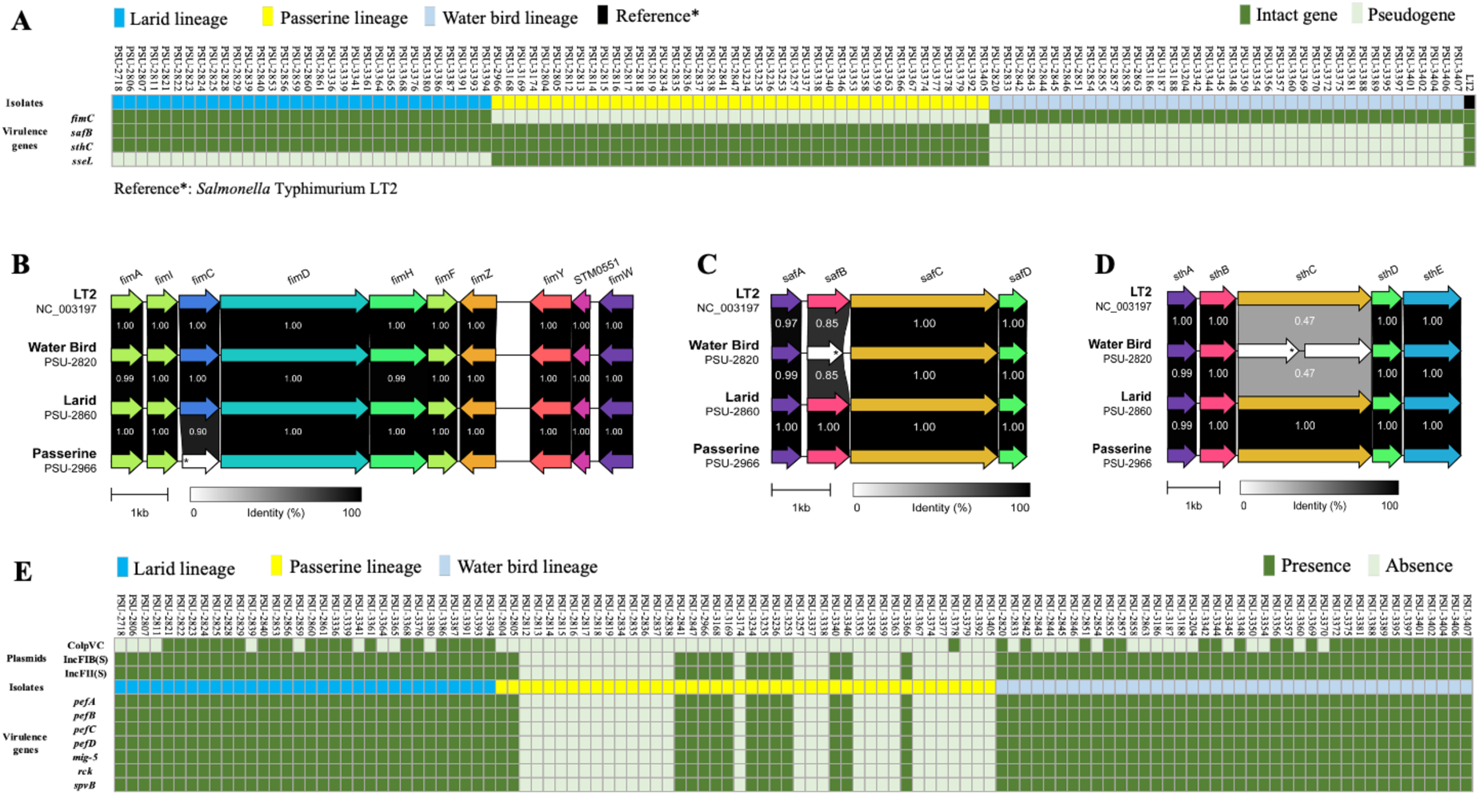
Chromosomal and plasmid-associated virulence gene signatures that differ among the larid, passerine, and water bird lineages. (A) Distribution of chromosomal virulence gene signatures (fimC, safB, sthC, and sseL) among the larid, passerine, and water bird lineages. (B-D) Organization of the fimbrial operons containing fimC, safB, and sthC in isolates from the larid, passerine, and water bird lineages, and reference strain S. Typhimurium LT2. (E) Distribution of plasmids and plasmid virulence genes among the larid, passerine, and water bird lineages. In the heatmaps (A) and (E): Light green = pseudogene in (A), and absence of the gene or plasmid in (E); Dark green = intact gene in (A), and presence of the gene or plasmid in (E); Blue = isolates from larid lineage; Yellow = isolates from passerine lineage; Light blue = isolates from water bird lineage; Dark = reference strain (*S*. Typhimurium LT2). Isolate names are represented on the x axis, virulence genes and plasmid names are represented on the y axis. Complete data used to generate the heatmaps is given in Table S3. In the fimbrial operons (B), (C), and (D), the pseudogenes are marked as white arrows, and the locations of indels in the pseudogenes are represented by asterisks.

Because of known concurrence, we conducted plasmid profiling to examine the correlation between certain virulence/resistance genes and plasmids in different wild bird lineages. As shown in **Fig. 4E**, four plasmids of the incompatibility groups ColpVC, IncFIB(S), IncFII(S), and IncI1 were present in these isolates. None of the plasmids carried antimicrobial resistance genes. However, three plasmids [IncI1, IncFIB(S), and IncFII(S)] carried virulence genes. IncI1 was present in a single isolate (PSU-2832); this plasmid carried the type IV pili genes (*pilUQRS*). IncFIB(S) and IncFII(S) carried virulence genes including *pefABCD* (<u>p</u>lasmid <u>e</u>ncoded <u>f</u>imbriae), *mig-5* (<u>m</u>acrophage induced <u>g</u>ene), *ric* (<u>r</u>esistance to <u>c</u>omplement <u>k</u>illing), and *spvB* (*<u>S</u>almonella* <u>p</u>lasmid <u>v</u>irulence). We found that 100% (72/72) of isolates from both water bird and larid lineages carried these two virulence plasmids, while only 33.3% (14/42) of isolates from the passerine lineage had these two plasmids. Isolates from passerine birds without the two plasmids also lacked the corresponding virulence genes (i.e., *pefABCD*, *mig-5*, *rck*, and *spvB*) (**Fig. 4E**).

## DISCUSSION

In this study, we explored the genetic diversity of 131 *S.* Typhimurium isolates collected from wild birds over four decades. Whole genome sequence phylogeny revealed three major lineages of *S.* Typhimurium largely defined by bird host taxonomic group; these lineages possessed certain STCTs and virulence gene signatures. Potential transmission of strains between the three lineages and non-avian host species was variable as was the extent of their geographic distribution. Our study also shows the wild bird *S.* Typhimurium lineages did not overlap with major lineages formed by domestic animal-sourced isolates. The addition of these wild bird isolates into a training dataset improved source prediction accuracy of a RF classifier among bovine, porcine, poultry, and wild birds.

Different host physiologies, habitats, and life histories may explain the divergence of *S.* Typhimurium lineages infecting different groups of wild birds. The three major lineages of *S.* Typhimurium in our study each corresponded to taxonomically disparate host groups. The water bird lineage was primarily associated with birds in the clade Aequornithes (e.g., cormorants, pelicans, herons), the larid lineage with birds in the order Charadriiformes (e.g., gulls, terns, plover), and the passerine lineage with birds in the order Passeriformes (e.g., finches, sparrows). Correspondence of these lineages to bird taxonomic group suggests a degree of host adaptation by *S.* Typhimurium infecting wild birds, similar to what has been described in strains infecting sea turtles (10) and pigeons (11). Some evidence suggests that certain *S.* Typhimurium strains maintained within passerine birds in the U.K. are host-adapted (12, 13). However, passerine-adapted *S.* Typhimurium strains in the U.K. belong to two definitive phage types (DT), i.e., DT40 and DT56(v) (the STCTs are 745, 7743, or 9520) (14, 15) and were apparently unrelated to the passerine isolates in this study, which mostly have the STCTs of 10, 11, and 34 (**Fig. 1A**). To the best of our knowledge, there are no readily available systematic studies reporting *S.* Typhimurium strains adapted to water birds or larids. Although the three lineages we describe herein were associated with particular bird groups at broad host taxonomic levels, we did not observe evidence of further specialization occurring at finer taxonomic scales. Specifically, *S.* Typhimurium strains isolated from the same host species did not form subclades within their major lineages (e.g., cormorant isolates did not form a subclade unique from pelican isolates within the water bird lineage). However, we sampled a relatively small number of wild bird hosts relative to the avian diversity that exists in North America. Specifically, our sampling was biased toward species which regularly experience outbreaks of salmonellosis such as cormorants, gulls, and finches (16).

Outbreaks may be more prevalent in these bird species because colonial nesting (cormorants, pelicans, herons, gulls, and terns) and congregation at feeding and water sites such as bird feeders and bird baths (songbirds) facilitate transmission. Thus, a greater diversity of host-adapted lineages of *S*. Typhimurium may be discovered by sampling additional avian groups that harbor cryptic infections.

Spatial patterns, including habitat partitioning by different bird hosts do not fully explain the *S.* Typhimurium phylogenetic patterns we observed. For example, cormorants, pelicans, and gulls often occupy the same habitats and occur in close proximity to one another. Yet, strains from these birds tended to fall into two distinct lineages based on bird phylogeny, suggesting factors such as host physiology may be the key factor driving *S.* Typhimurium host adaptation. Notably, the passerine lineage is sister to the larid lineage, and both are more distantly related to the water bird lineage despite water birds and larids being much more closely related than either group is to the passerines (17). The Bayesian phylogenetic inference indicates the three lineages formed sometime after 1900. This implies that host adaptation occurred well after the divergence of the avian host groups. Similarly, a lineage of *S*. Typhimurium adapted to sea turtles across the Pacific Ocean evolved from a MRCA only a few decades ago (10). Taken together, these findings suggest that spillover of *S*. Typhimurium into wildlife and subsequent host adaptation may be a relatively recent phenomenon, likely driven by anthropogenic influences.

While the presence of three major lineages suggested host adaptation by bird-infecting strains of *S.* Typhimurium, there was evidence that strains within a lineage could infect bird species outside of its typical host group. Infections in aberrant hosts usually corresponded with hosts from different groups that share similar environments to one another. For example, *S.* Typhimurium strains in the water bird lineage were occasionally isolated from other birds (i.e., terns, gulls, grebe, and goose) that share aquatic habitats with cormorants, pelicans, and herons. Interestingly, strains of *S.* Typhimurium isolated from raptors (e.g., eagles, hawks, and owls) were phylogenetically dispersed. The diversity of strains harbored by raptors may be due to acquisition of *S.* Typhimurium through consumption of other birds and animals that may be infected with the bacterium rather than possessing their own unique strains.

While *S.* Typhimurium strains clustered by host species, there was little evidence of geographic patterns based on collection locations within the U.S. (**Fig. 1A**). This may be explained by the high mobility of wild birds, which travel long distances and across political boundaries. Isolates from the three major lineages did, however, cluster with isolates from different global regions in the PD database. Strains from the larid lineage were genetically similar to isolates in the database originating from South America, Oceania, and Europe. This is consistent with the long-distance migratory capacity for terns and gulls which makes them capable of spreading certain strain of *S.* Typhimurium across the world. Isolates from the water bird lineage exhibited a similar geographic pattern. In contrast, isolates from the passerine lineage did not closely match isolates outside of North America, which is consistent with the more typical intracontinental migratory patterns for the passerine species. It should be noted that the PD database is heavily skewed towards isolates from the U.S. Isolates from other geographic regions are underrepresented since many countries do not sequence isolates or actively submit data to the NCBI PD database for surveillance purposes. Additional sampling in other parts of the world may reveal that all three major lineages are more widely distributed.

Host adaptation of *S.* Typhimurium in various groups of wild birds may affect their transmission among different hosts or sources. For example, isolates from the water bird and larid lineages clustered with isolates from aquatic animals (fish/shellfish), which is understandable given their overlap in habitats. However, no isolates from aquatic animals were closely related with isolates from the passerine lineage. Instead, the passerine lineage was more likely to contain *S.* Typhimurium strains isolated from horses and cats. Studies have reported that cats can acquire *S.* Typhimurium by catching infected songbirds or spending time around bird feeders (18, 19). However, few studies explored the potential spread of *S.* Typhimurium between wild birds and horses, which may be exposed to strains of the passerine lineage in open pastures or stables that are also occupied by avian species such as finches and house sparrows.

Interestingly, isolates from the three wild bird lineages were rarely related to isolates from livestock and poultry, but were frequently associated with clinical isolates from human beings (**Fig. 2** and Table S2). Our addition of isolates to the database will facilitate attribution of *S.* Typhimurium strains from wild birds recovered from other hosts or substrates to the appropriate source. Although the exact transmission route and direction are difficult to determine, studies such as this are important in raising awareness of wild birds as a reservoir of *S.* Typhimurium that may contribute to preventable cases of human salmonellosis. For example, several studies have reported the concurrence of *S.* Typhimurium outbreaks in both passerine birds and humans in the U.K., U.S., and Sweden during late winter or early spring (6, 19, 20), and cats are considered to be a possible intermediate vehicle for *S.* Typhimurium transmission between passerine birds and human beings during outbreaks (19, 21, 22). This is consistent with our observation that passerine bird isolates cluster with isolates from cats in the PD database (**Fig. 2C**). In 2021, an *S.* Typhimurium outbreak that caused at least 29 illnesses and 14 hospitalizations in the U.S. was traced back to isolates from songbirds (7). In fact, clinical isolates from this outbreak formed a SNP cluster with passerine isolates in our study (**Fig. 5**; three pine siskin isolates: PSU-3338, PSU-3367, and PSU-3392; two redpoll isolates: PSU-3337; and one crossbill isolate: PSU-3353, highlighted in red). These bird isolates belonged to STCT10 and STCT11. Our passerine isolates differed from clinical isolates by <10 SNPs, suggesting a recent common ancestor (23). None of our passerine isolates were collected earlier than the clinical isolates from the 2021 outbreak (**Fig. 5**), and although we cannot say with certainty, it is plausible that strains transmitted from passerine birds to humans were responsible for the outbreak. Humans may get infected via direct contact with songbirds, contaminated bird feeders, or infected cats; or there may be additional unknown sources not represented in the PD database. To further elucidate routes of transmission, collecting contemporaneous *S*. Typhimurium isolates from humans, wild birds, and environmental samples through outbreak investigations is necessary. Such efforts offer insights to guide preventative measures to reduce the health risk for future outbreaks.

**FIG 5.**
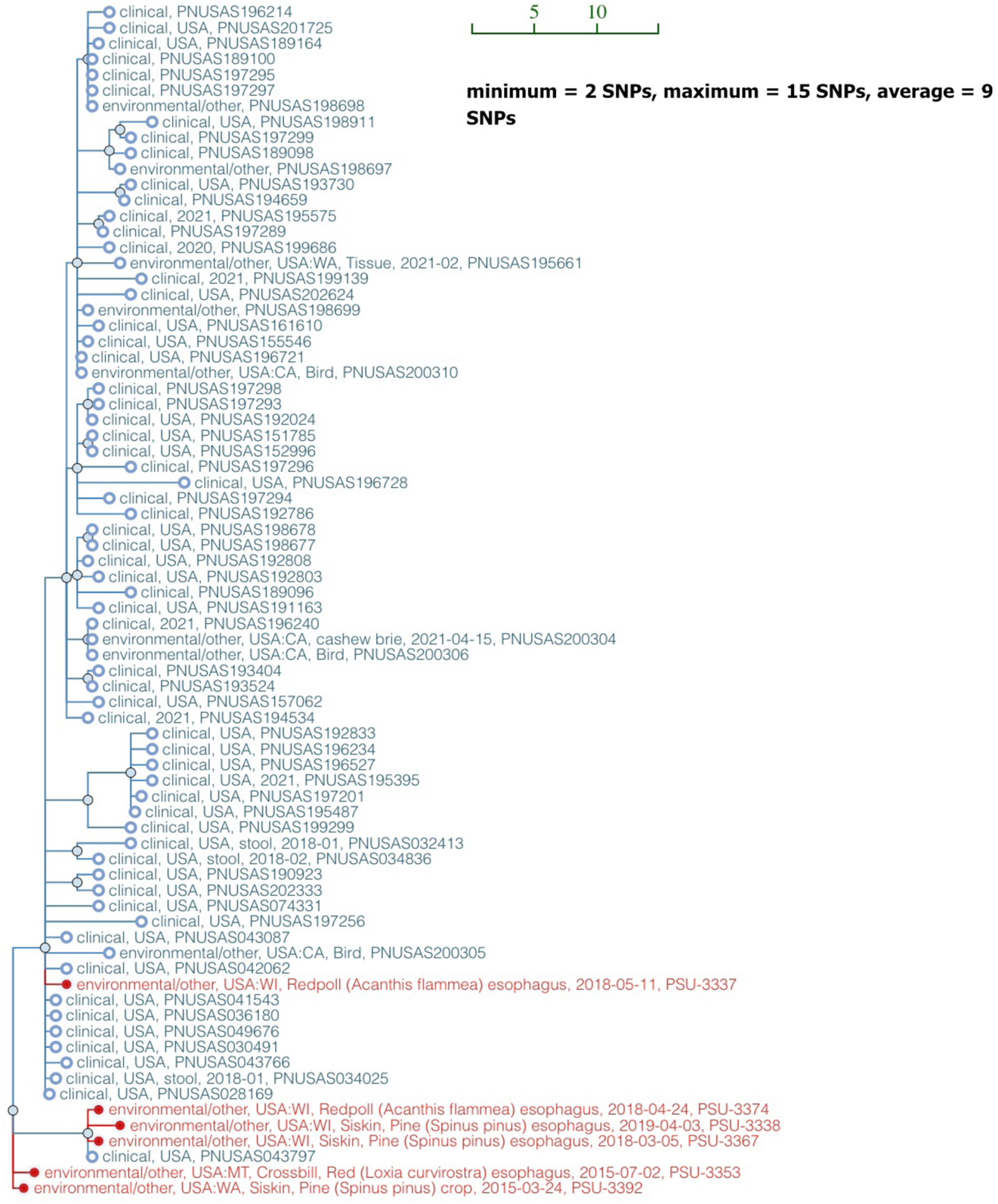
Close genetic relatedness between passerine isolates from this study and clinical isolates from a 2021 salmonellosis outbreak in the U.S. Single nucleotide polymorphism (SNP) tree is automatically generated in the NCBI Pathogen Detection database when sequencing data are submitted to the database (SNP cluster and accession link: <u>PDS000013825.81</u>). The scale bar represents the unit SNP distance. The average SNP distance for all isolates in the tree is nine. *S*. Typhimurium isolates in the SNP tree from this study are highlighted in red.

Inclusion of the wild bird isolates in the PD database not only facilitates outbreak investigation, but also improves source attribution. The RF classifier in this study performed much better to predict isolates from wild birds with the addition of our 131 isolates to a training dataset from porcine (*n* = 338), bovine (*n* = 195), poultry (*n* = 440), and wild birds (*n* = 68) (**Table 1**). The significant increase in wild bird isolates from 68 to 199 may contribute to the improved prediction accuracy for wild bird sources. It should be noted, however, that prediction accuracy not only depends on sampling intensity but also presence of representative isolates from different sources.

Even though the count of isolates from bovine sources (*n* = 195) is almost equal to that from wild birds (*n* = 199), prediction accuracy for bovine sources (62.56%) is lower than that for wild bird sources (83.42%). The high prediction accuracy for wild birds may also be attributed to consistent presence of genetic signatures such as pseudogene accumulation in fimbrial genes which may help discriminate wild bird isolates from other sourced isolates. This is consistent with the observation that wild bird isolates form three unique lineages, while bovine isolates are more phylogenetically dispersed in the SNP phylogenetic tree (**Fig. 3**). Overall, an improvement in prediction accuracy is crucial to help identify root sources of *S.* Typhimurium outbreaks, especially when the outbreak is linked to nonfood sources.

It is well documented that genome degradation contributes to *Salmonella* host adaptation (24, 25, 26, 27, 28, 29). Our study shows pseudogene accumulation in specific chromosomal fimbrial genes in certain wild bird *S.* Typhimurium lineages (i.e., *fimC* in passerine lineage, *safB* and *sthC* in water bird lineage). The inactivation of these genes due to frameshift and premature stop codons may help *S.* Typhimurium adapt to a particular bird host. Furthermore, passerine bird isolates tended to lose the plasmid carrying virulence genes (i.e., *pefABCD*, *mig-5*, *rck*, and *spvB*). These virulence genes are important for *S.* Typhimurium survival and replication in human and mouse macrophages (30, 31, 32), which may indicate isolates from the passerine lineage are more host-adapted to birds than those from water bird and larid lineages.

To conclude, the findings in this study highlight the importance of wild birds as a reservoir for *S.* Typhimurium. Source attribution can be more precise if we include more representative wild bird isolates into *S.* Typhimurium databases. In addition, the distinct lineages defined by bird type, together with the presence of virulence gene signatures, imply host adaptation of *S.* Typhimurium to wild birds. Considering wild birds may spread *S.* Typhimurium in a very different manner than domestic animals, alternative management strategies will be required to prevent their transmission to human beings. While wild birds serve as potential reservoirs of *S.* Typhimurium, these species also provide critical ecological and economic services to humans and are of great cultural significance. Activities aimed at preventing zoonotic transmission should focus on strategies that benefit both wild birds and humans alike.

## MATERIALS AND METHODS

### Bacterial strains

*Salmonella* Typhimurium strains (*n* = 131) from wild birds were isolated by the U.S. Geological Survey - National Wildlife Health Center between 1978 and 2019 from 30 states in the U.S. and stored in cryogenic vials at −80 °C. All strains were confirmed as *S.* Typhimurium by traditional *Salmonella* serotyping and reconfirmed by SeqSero2 v1.2.1 (33) from newly generated whole genome sequencing data (see below).

### DNA extraction and whole genome sequencing

For DNA extraction, each isolate was streaked onto xylose lysine deoxycholate (XLD) agar plates and incubated for 18 h at 37 °C. A single colony was then picked, transferred to Luria-Bertani broth (LB) and cultured overnight at 37 °C with continuous agitation (250 rpm). Genomic DNA was extracted using the Qiagen DNeasy^®^ Blood & Tissue kit (Qiagen, Valencia, CA) following the manufacturer’s instructions. Genomic DNA purity was confirmed via an A260/A280 measurement (target ≥1.8) using NanoDrop™ One (Thermo Scientific™, DE), and DNA concentration was quantified using Qubit^®^ 3.0 (Thermo Fisher Scientific Inc., MA) fluorometer.

Isolated genomic DNA was stored at −20 °C before use. For whole genome sequencing (WGS), genomic DNA was adjusted to 0.2 ng/μL. DNA library was then prepared using the Nextera XT DNA Library Prep Kit (Illumina Inc., San Diego, CA). The resulting library was normalized using quantitation-based procedure and pooled together at equal volume. The pooled library (600 μL) was denatured and sequenced on an Illumina MiSeq sequencer (Illumina Inc., San Diego, CA) using a MiSeq reagent v3 kit, with 500 (2 x 250) cycles.

### Quality assessment for raw reads

After sequencing, the quality of Illumina paired-end reads of all isolates were assessed using the MicroRunQC workflow in GalaxyTrakr v2 (34). Raw reads passing quality control thresholds (i.e., average coverage >30, mean quality score >30, number of contigs <400, total assembly length between 4.4-5.l Mb) were submitted to the NCBI under BioProject <u>PRJNA357723</u>. Metadata information and Sequence Read Archive (SRA) accession numbers of these isolates are listed in **Table S1.**

### Phylogenetic analysis

For genetic relatedness comparison between the 131 isolates, raw reads were uploaded to Enterobase (35). *S.* Typhimurium strain LT2 (RefSeq NC_003197.1) served as the reference genome and a maximum-likelihood (ML) phylogenetic tree was created using the SNP project in Enterobase based on 5,839 SNPs in the core genomic regions of the 131 wild bird *S.* Typhimurium isolates. The ML phylogenetic tree was re-constructed by MEGA X v10.1.8 using the Tamura-Nei model and 500 bootstrap replicates (36). The SNP phylogenetic tree was visualized and annotated using the Interactive Tree of Life (iTOL v6; https://itol.embl.de). CRISPR arrays were identified using CRISPRviz (37). Spacers were aligned and unique arrays were given a unique allelic identifier as described by Shariat et al (38). *S.* Typhimurium CRISPR Type (STCT) was then determined by the unique combination of CRISPR1 and CRISPR2 alleles. In addition, sequence type (ST) of these *S.* Typhimurium isolates was identified using 7-gene (*aroC*, *dnaN*, *hemD*, *hisD*, *purE*, *sucA* and *thrA*) multilocus sequence typing (MLST) at Enterobase (35). STCTs and STs were annotated in the SNP phylogenetic tree.

### Bayesian phylogenetic inference

The evolutionary history of the 131 isolates was inferred by Bayesian phylogenetic analysis. Snippy (Galaxy v4.5.0) (https://github.com/tseemann/snippy) was used to generate a full alignment and find SNPs between the reference genome LT2 (RefSeq NC_003197.1) and the genomes of wild bird isolates, and Snippy-core (Galaxy v4.5.0) (https://github.com/tseemann/snippy) was used to convert the Snippy outputs into a core SNP alignment. The resultant multiple sequence alignment (5,251 variant sites) was used to construct a time-scale Bayesian phylogenetic tree by BEAUti v2.6.5 and BEAST2 v2.6.5 (39). The parameters were set as followings: Prior assumption-coalescent Bayesian skyline; Clock model-relaxed clock log normal; Markov chain Monte Carlo (MCMC): chain length-100 million, storing every 1,000 generations. A maximum clade credibility tree was created using TreeAnnotator v2.6.4 with burnin percentage of 10% and node height option of median height (39). Finally, the tree was visualized using FigTree v1.4.4 (https://github.com/rambaut/figtree/releases).

### Genetic relatedness between wild bird isolates and other isolates

The genetic relatedness between wild bird isolates and other isolates from environmental sources, food, or human hosts were inferred by using the NCBI Pathogen Detection https://www.ncbi.nlm.nih.gov/pathogens/. After uploading the raw reads of the 131 isolates to NCBI, NCBI Pathogen Detection assembled, annotated, and clustered the newly sequenced isolate genomes to other closely related isolates in the database. Clustering involved two steps: First, related isolates were clustered based on whole genome MLST (wgMLST) scheme of *Salmonella* with a 25-allele cut-off; Once wgMLST clusters were created, SNPs are called by aligning assemblies against a reference genome chosen from each wgMLST cluster of closely related isolates, and SNP clusters and phylogenetic trees were inferred. A SNP cluster was classified as a cluster of isolates where each isolate was less than or equal to 50 SNPs distant from other members of the cluster (40). Individual phylogenetic trees for each SNP cluster, together with the metadata information of isolates in the same cluster (**Table S2**), were used to examine relationship between wild bird isolates and other isolates.

### Random Forest-based source attribution

A machine learning Random Forest (RF) classifier was used for source attribution of *S.* Typhimurium to bovine, porcine, poultry and wild bird sources. The RF classifier was built using the method described by Zhang et al (9). Briefly, a set of 3,102 genetic features, including 1,850 SNPs, 147 indels, and 1,105 accessory gene, were used for the RF classifier. The randomForest package (v4.6-12) of R was used to build the RF classifier with the “ntree” argument set to be 1,000 and other parameters being default. The original training data (195 bovine isolates, 338 porcine isolates, 440 poultry isolates, and 68 wild bird isolates) and the training data with the addition of the 131 isolates from wild birds were applied for classifier training. Tables of confusion were generated to describe the performance of the classifier. To explain the prediction performance of the classifier after adding our bird isolates into the training dataset, a maximum-likelihood tree based on core genome alignment of 1,604 *S.* Typhimurium isolates from different sources was constructed using Parsnp v1.5.0 (41). The 1,604 *S.* Typhimurium isolates included 171 human clinical isolates, 83 food isolates, 178 isolates from miscellaneous sources, and 1,172 isolates used for the RF classifier training (i.e., 195 bovine isolates, 338 porcine isolates, 440 poultry isolates, and 199 wild bird isolates in which 131 isolates were from this study).

### AMR, virulence, and plasmid profiling

Raw reads of each isolate were *de novo* assembled using Shovill with Trimmomatic on (Galaxy v1.0.4) (42). To identify the antimicrobial resistance (AMR) genes present in the isolates, each draft genome assembly was searched against ResFinder database and CARD database using ResFinder 4.1 (43) and Resistance Gene Identifier (RGI) v5.2.0 (44). AMR genes that passed the default threshold values for ResFinder (≥90% nucleotide identity and ≥60% coverage) and RGI (≥95% nucleotide identity) were considered to be present in the isolate. Virulence factors were predicted by aligning the draft genome assembly for each isolate against the Virulence Factor database using VFanalyzer (45). Virulence factors that passed the default threshold values for VFanalyzer (≥90% amino acid identity and ≥50% coverage) were considered to be present in the isolate. If a virulence factor was absent in the isolate identified by VFanalyzer, the virulence gene encoding the specific virulence factor was then compared to its reference gene from strain *S.* Typhimurium LT2 by BLAST to confirm the mutation type (e.g., insertion, deletion, or substitution). Plasmid replicon sequences were identified by comparing the draft genome assembly against the PlasmidFinder database using PlasmidFinder 2.1 with default search settings of ≥95% nucleotide identity and ≥60% coverage (46).

### Data availability

Sequence data of the 131 *S.* Typhimurium strains are deposited in the NCBI Sequence Read Archive (https://www.ncbi.nlm.nih.gov/sra) under BioProject <u>PRJNA357723</u>. Accession numbers are available in **Table S1**. Strains that form SNP clusters with wild bird isolates examined in this study are provided in **Table S2** and publicly available in the NCBI Pathogen Detection (https://www.ncbi.nlm.nih.gov/pathogens/).

## ACKNOWLEDGEMENTS

We thank Drs. Heather Tate and Patrick McDermott for editing this manuscript prior to publication.

This work is supported by the U.S. Food and Drug Administration under Grant No. 1 U19 FD007114-01 (to E.G.D.), U.S. Department of Agriculture under Grant No. PEN4522 (to E.G.D.), and Penn State College of Agricultural Sciences.

The views expressed in this article are those of the authors and do not necessarily reflect the official policy of the Department of Health and Human Services, the U.S. Food and Drug Administration, or the U.S. Government. Any use of trade, firm, or product names is for descriptive purposes only and does not imply endorsement by the U.S. Government.

## Author contributions

Y.F. whole genome sequenced the wild bird isolates, designed the study, performed the majority bioinformatics analysis of the data, interpreted the data, and wrote the draft manuscript. N.M.M., J.M.L., D.S.B., B.B-Z., C.A.W., and E.G.D. assisted in collection of the isolates and contributed to interpretation of data and manuscript revision. S.L. and X.D. participated in Random Forest-based source attribution, interpretation of data, and manuscript revision. J.C.S. and N.W.S. participated in *S.* Typhimurium CRISPR typing, interpretation of data, and manuscript revision. E.M.N contributed to identification of virulence gene mutation, interpretation of data, and manuscript revision. E.G.D. additionally designed and supervised the study.

The authors declare no competing interests.

## Legends for Supplementary Materials

FIG S1 Comparison of SseL from different wild bird isolates. Multiple sequence alignment of SseL amino acid sequences from the larid, passerine, and water bird lineages. SseL from *S.* Typhimurium LT2 served as a reference.

Table S1 Metadata information (isolate name, SRA accession number, isolation year, state, bird host, CRISPR type, and 7-gene MLST sequence type) of the 131 *S.* Typhimurium isolates from wild birds collected between 1978-2019 in 30 U.S. states.

Table S2 SNP clusters generated by the NCBI Pathogen Detection when relating the isolates from larid, passerine, and water bird lineages to other isolates in the database.

Table S3 *In silico* antimicrobial resistance, virulence, and plasmid profiles of the 131 *S.* Typhimurium isolates from wild birds collected between 1978-2019 in 30 U.S. states.

