## Supplemental Figure S1 for "*Salmonella enterica* serovar Typhimurium from Wild Birds in the United States Represent Distinct Lineages Defined by Bird Type"

```
LT2_sseL            1 VSDEALTLLFSAVENGDQNCIDLLCNLALRNDDLGHRVEKFLFDLFSGKRTGSSDIDKKI
passerine_sseL      1 VSDEALTLLFSAVENGDQNCIDLLCNLALRNDDLGHRVEKFLFDLFSGKRTGSSDIDKKI
larid_sseL          1 VSDEALTLLFSAVENGDQNCIDLLCNLALRNDDLGHRVEKFLFDLFSGKRTGSSDIDKKI
water_sseL          1 VSDEALTLLFSAVENGDQNCIDLLCNLALRNDDLGHRVEKFLFDLFSGKRTGSSDIDKKI
consensus           1 ************************************************************

LT2_sseL           61 NQACLVLHQIANNDITKDNTEWKKLHAPSRLLYMAGSATTDLSKKIGIAHKIMGDQFAQT
passerine_sseL     61 NQACLVLHQIANNDITKDNTEWKKLHAPSRLLYMAGSATTDLSKKIGIAHKIMGDQFAQT
larid_sseL         61 NQACLVLHQIANNDITKDNTEWKKLHAPSRLLYMAGSATTDLSKKIGIAHKIMGDQFAQT
water_sseL         61 NQACLVLHQIANNDITKDNTEWKKLHAPSRLLYMAGSATTDLSKKIGIAHKIMGDQFAQT
consensus          61 ************************************************************

LT2_sseL          121 DQEQVGVENLWCGARMLSSDELAAATQGLVQESPLLSVNYPIGLIHPTTKENI-------
passerine_sseL    121 DQEQVGVENLWCGARMLSSDELAAATQGLVQESPLLSVNYPIGLIHPTTKENI-------
larid_sseL        121 DQEQVGVENLWCGARMLSSDELAAATQGLVQESPLLSVNYPIGLIHPTTKENI-------
water_sseL        121 DQEQVGVENLWCGARMLSSDELAAATQGLVQESPLHLGKLSYWADSSYHQRKYIKHSAT*
consensus         121 ***********************************..................

LT2_sseL          174 --LSTQLLEKIAQSGL---SHNEVFLVNTGDHWLLCLFYKLAEKIKCLIFNTYYDLNENT
passerine_sseL    174 --LSTQLLEKIAQSGL---SHNEVFLVNTGDHWLLCLFYKLAEKIKCLIFNTYYDLNENT
larid_sseL        174 --LSTQLLEKIAQSGL---SHNEVFLVNTGDHWLLCLFYKLAEKIKCLIFNTYYDLNENT
water_sseL        180 KDCSIRIISQ*SLPGKYRRSLASLFIL*TCR------KNKMPYI*YLL*FK*KY*ARDYR
consensus         181   .*..........*.   *....*...*...........*......*.*...*......

LT2_sseL          229 KQ-EIIEAAKIAGISESDE------VNFIEMNLQ---NNVPNGCGLFCYHTIQLLSN---
passerine_sseL    229 KQ-EIIEAAKIAGISESDE------VNFIEMNLQ---NNVPNGCGLFCYHTIQLLSN---
larid_sseL        229 KQ-EIIEAAKIAGISESDE------VNFIEMNLQ---NNVPNGCGLFCYHTIQLLSN---
water_sseL        228 SSKNCRHIRKR*G*FY*NEFTEQCTQRLWSILLPYNSTLIECRTKRSCYHTTRICGKFLN
consensus         241 .. ......*..*.....*      .......*.   ..........****......

LT2_sseL          276 --------AGQNDPATTLREFAENFLTLSVEEQALFNTQTRRQIYEYSLQ*
passerine_sseL    276 --------AGQNDPATTLREFAENFLTLSVEEQALFNTQTRRQIYEYSLQ*
larid_sseL        276 --------AGQNDPATTLR*FAENFLTLSVEEQALFNTQTRRQIYEYSLQ*
water_sseL        285 AFSRGTSTI*HPNPAANI*IQS------PV--------------------
consensus         301         .....**..............*....................
```
